# Coiled Coil-Mediated Assembly of an Icosahedral Protein Cage with Extremely High Thermal and Chemical Stability

**DOI:** 10.1101/316331

**Authors:** Ajitha S. Cristie-David, Junjie Chen, Derek B. Nowak, Sung I. Park, Mark M. Banaszak Holl, Min Su, E. Neil G. Marsh

## Abstract

The organization of protein molecules into higher-order nanoscale architectures is ubiquitous in Nature and represents an important goal in synthetic biology. Here we describe the symmetry-directed design of a hollow protein cage with dimensions similar to those of many icosahedral viruses. The cage was constructed based on icosahedral symmetry by genetically fusing a trimeric protein (TriEst) to a small pentameric de novo-designed coiled coil domain, separated by a flexible oligo-glycine linker sequence. Screening a small library of designs in which the linker length varied from 2 to 12 residues identified a construct containing 8 glycine residues (Ico8) that formed well-defined cages. Characterization by dynamic light scattering, negative stain and cryo EM, and by atomic force and IR-photo-induced force microscopy established that Ico8 assembles into a flexible hollow cage with comprising 60-subunits with overall icosahedral geometry. Unexpectedly, the cages were found to encapsulate DNA, even though neither protein component binds nucleic acids on its own. Notably, the cages formed by Ico8 proved to be extremely stable towards thermal and chemical denaturation: whereas TriEst was unfolded by heating (Tm ~75 °C) or denatured by 1.5 M guanidine hydrochloride, the Ico8 cages remained folded even at 120 °C or in 8 M guanidine hydrochloride. The encapsulation of DNA and increased stability of the cages are new properties that emerge from the higher order structure of the protein cage, rather than being intrinsic to the components from which it is constructed.

## Introduction

Self-assembling proteins are attractive as building blocks for “smart” biomaterials that can potentially exploit the rich structural and functional properties of proteins (1-4). This has spurred efforts to design new self-assembling protein structures by applying symmetry principles to construct geometric cages and extended lattices (5-10). Successful designs have utilized computational approaches to design new interfaces between protein subunits to facilitate assembly (3, 11-13), or the fusion of two oligomeric proteins through a rigid alpha-helical linker (14). Using these strategies, tetrahedral, cubic (14), octahedral (12, 13), dodecahedral and icosahedral (3, 11) protein cages as well as 2- and 3-dimensional (15, 16) protein lattices have been constructed. However, these approaches require that the angle between the two rotational symmetry axes that defines the geometry of the assembly needs to precisely specified for the protein subunits to assemble correctly. This has proved technically challenging to accomplish.

Our laboratory has developed an alternative design strategy, in which protein assembly is mediated by short, parallel, coiled coil domains that are genetically fused to the larger building block protein through a short, flexible spacer sequence (17-19). This approach, greatly simplifies the design process because angle between the symmetry axes does not need to specified. By using combinations of symmetry axes that are unique to the geometry of the desired protein cage, it is possible to assemble well-defined protein cages using symmetry considerations alone. Thus, we have recently described the construction of octahedral (19) and tetrahedral (17) cages by combining a C_3_-symmetric building block protein with either a C_4_-symmetic or C_3_-symmetric coiled coil assembly domain, respectively. This approach is inherently modular, as many coiled designs are available and there are few constraints on the choice of protein building block; it is also far less computationally intensive than protein interface re-design.

Here, we demonstrate the extension of this approach to the design of an icosahedral cage. As the largest Platonic solid, icosahedral protein cages can encapsulate large volumes and as such are often used to in Nature for packaging and transport (20). Large, robust and customizable protein cages should be useful in synthetic biology (1, 21), vaccine design (22) and targeted drug delivery (23). However, icosahedral cages remain challenging to construct, as they require the coordinated assembly of 60 protein subunits to achieve the desired icosahedral geometry.

For the combination of C_3_ and C_5_ symmetric proteins necessary to specify an icosahedral cage, we selected a trimeric esterase (TriEst) as the protein building block and a well-characterized pentameric parallel coiled-coil domain. These were linked together through a flexible oligo-Gly sequence. Characterization of the resulting protein assemblies revealed that they form hollow cages, with features consistent with the intended 60-subunit icosahedral geometry. Notably, assembly of the trimeric esterase into a cage resulted in a remarkable increase in both its thermal stability and resistance to chemical denaturation by guanidium hydrochloride (GuHCl) and extremes of pH.

## Results

To design a self-assembling icosahedral protein cage, we followed the approach previously taken to design tetrahedral and octahedral cages. For the C_3_-symmetric building block we used the same trimeric esterase (PDB ID: 1ZOI) as in our previous studies to facilitate comparisons with our earlier designs and assess the generality of the design strategy (24). To provide the C_5_ component necessary for icosahedral geometry we selected a de novo-designed, parallel, pentameric coiled coil (25) (PDB ID: 4PN8); this component was modified by mutating W13Q to alleviate problems with aggregation, as described previously (18).

### Screening and Optimization of Assembly

Synthetic genes were constructed to encode proteins in which the C-terminus of TriEst was linked to the N-terminus of the coiled coil through a flexible sequence containing increasing numbers of Gly residues. (Fig. 1 & Table S1). The number of Gly residues were increased two at a time to give constructs containing between 2 and 12 Gly, herein referred to as Ico2 through Ico12 respectively, with each additional di-Gly unit increasing by ~ 5 Å the distance between the C_3_ and C_5_ domains. These constructs were expressed and purified by standard procedures as described in the methods section.

**Figure 1.**
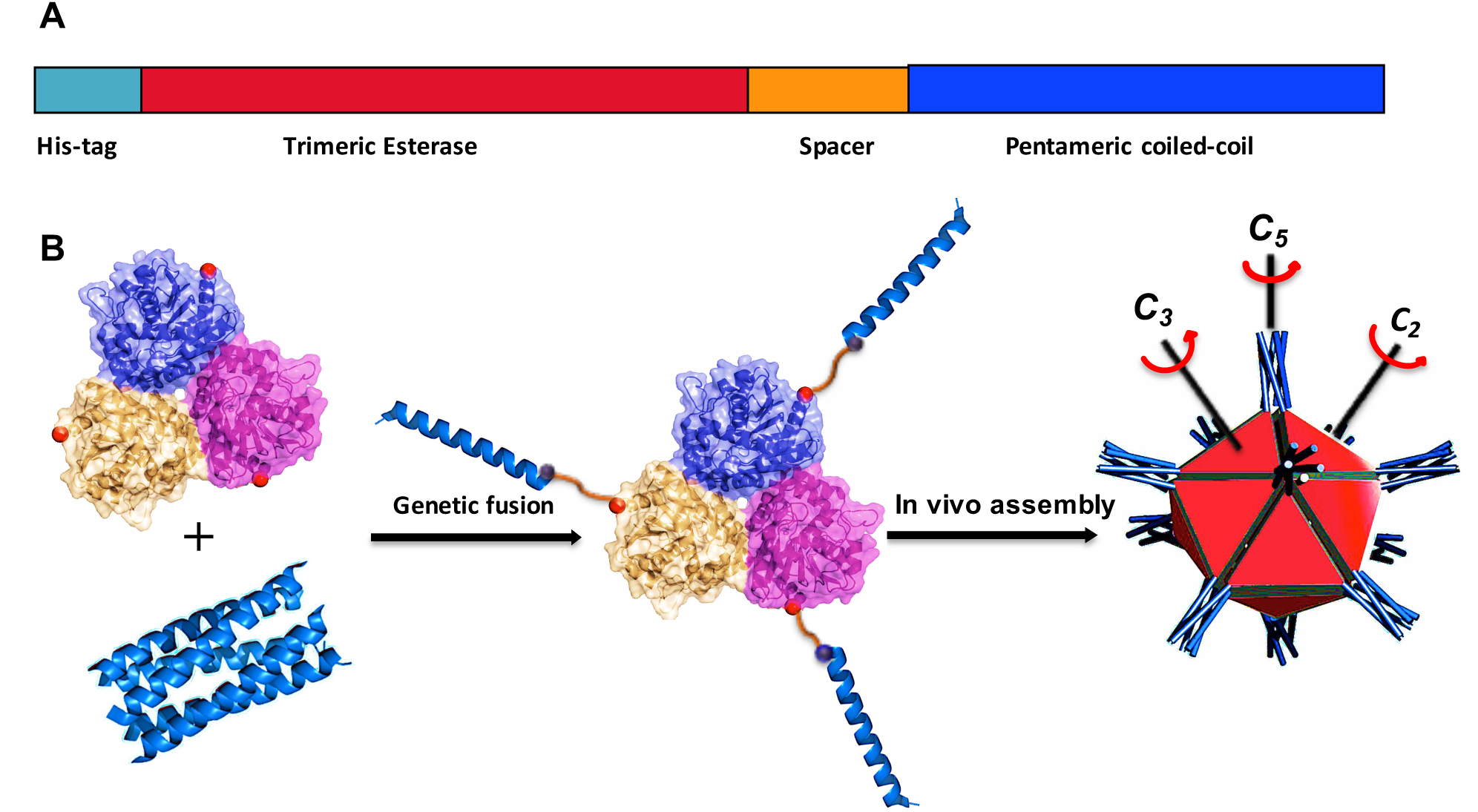
Design scheme for a self-assembling icosahedral protein cage. **A)** The Ico8 fusion protein comprises a trimeric esterase an oligo-Gly linker sequence and a pentameric coiled coil. **B)** Structures of the trimeric esterase and pentameric coiled coil with a schematic illustration of the assembly of the fusion protein into an icosahedral cage.

Preliminary characterization of Ico2 and Ico4 indicated that the proteins were unstable and rapidly precipitated, presumable because the linker length was too short for the protein domains to fold properly. The other constructs were sufficiently stable to be characterized by SEC and negative stain EM. Ico6, Ico10 and Ico12 each formed a heterogeneous mixture of assembled proteins, aggregated material and unassembled trimers (Fig. S1). However, Ico8 assembled into spherically shaped assemblies that were of approximately the size expected for a 60-subunit protein cage (estimated to be ~ 27 nm in diameter), as determined by size exclusion column chromatography (SEC), dynamic light scattering and negative stain EM (Fig 2). The size distribution, estimated from analysis of negative stain EM images of the particles was quite broad, ranging from ~24 – 30 nm in diameter. These findings were consistent with the flexible nature of the design, as the oligo-Gly spacer sequence has the potential to span a length of ~ 20 Å. However, we don’t rule out the possibility that some particles may have misassembled and contain either fewer or greater than the intended 60 protein subunits. Higher resolution images were obtained using scanning transmission electron microscopy (STEM) (Fig. 2C). These images revealed hexagonal features in many of the particles, particularly apparent in the Fourier-filtered images, which is consistent with the intended icosahedral geometry of the protein cages.

**Figure 2.**
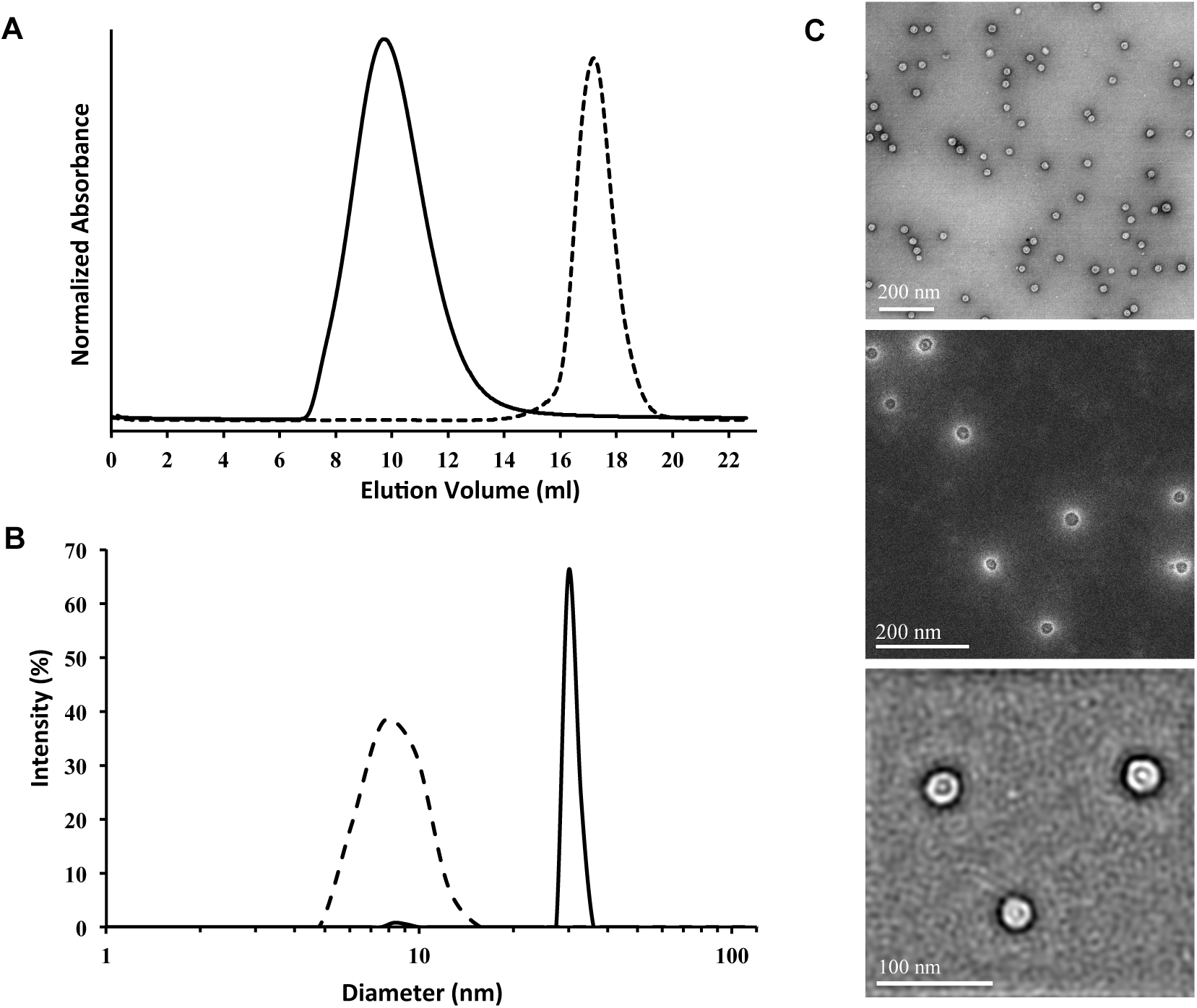
Characterization of the protein cages formed by Ico8. **A)** Size and homogeneity of Ico8 assessed by size exclusion chromatography: solid line, Ico8; dashed line TriEst; **B)** Number-averaged diameter distribution of Ico8 cages measured dynamic light scattering: solid line, Ico8; dashed line TriEst **C)** *Top*: Conventional negative stain EM of particles formed by Ico8; *middle*: high-angle annular dark-field scanning EM of Ico8 particles; *bottom*, bright field Fourier-filtered scanning EM images reveal hexagonal features associated with cages.

Based on these initial studies, the protein cages formed by Ico8 were selected for further detailed characterization. Although Ico8 contained an N-terminal His tag, the protein bound only weakly to Ni-NTA resin used in the initial purification. We therefore took advantage of the large size and heat stability of the assembled protein (discussed later) to purify Ico8. This was achieved by heating the cell lysate at 70 °C for ~ 75 min to precipitate most *E. coli* proteins, which were removed by centrifugation. After dialysis, the cages were purified further by two rounds of SEC. This procedure produced electrophoretically pure material (Fig. S1) that was used for further characterization.

### Atomic force microscopy of Ico8

To obtain further information on the size of the cages formed by Ico8, we imaged the cages using atomic force microscopy (AFM, Fig. 3A), which has the advantage of directly measuring the height and volume of particles. Particles were spin-coated onto a mica substrate as described in the Methods section and imaged using a carbon nanotube (CNT) tip in AC mode. ~1100 particles were imaged and subjected to automated particle analysis and classification (Fig 3B, Fig S2). Consistent with their flexible, hollow nature, the cages flattened onto the freshly cleaved mica surface resulting in an average particle height of 3.6 ± 1.1 nm. To compare to the theoretically expected volume and the experimental volume obtained from cryo-electron microscopy (cryo-EM), the measured particle volumes were used to calculate the diameter of an ideal icosahedral geometry. This yielded an average diameter of 24.9 ± 7.0 nm for the Ico8 cages, with the distribution of particle sizes being in good agreement with that measured by both cryo-EM and negative stain EM (*vida infra*).

**Figure 3.**
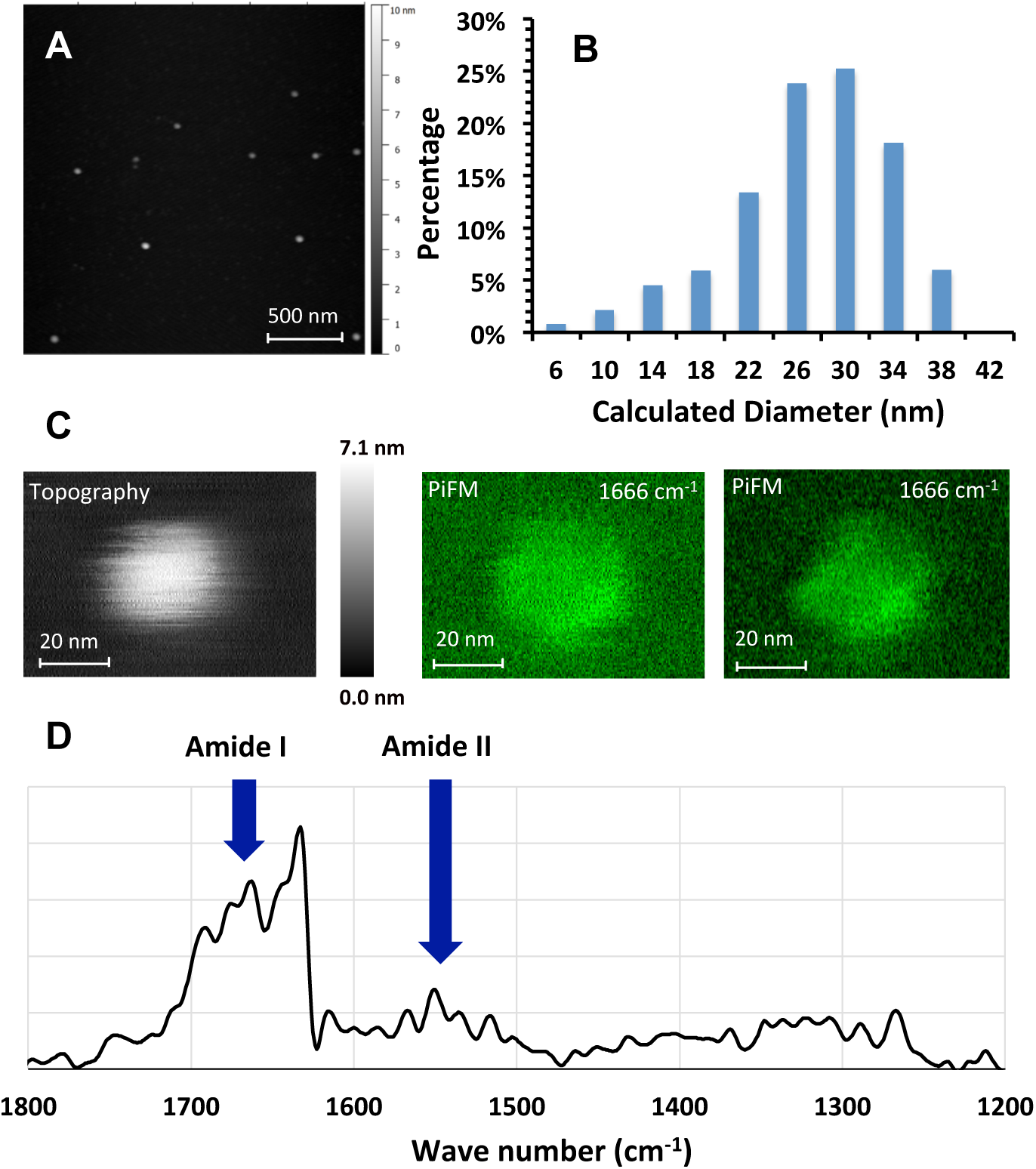
Atomic force microscopy and IR-photo-induced force microscopy of Ico8 cages. **A)** Representative field of view of Ico8 particles imaged by AFM. **B)** Distribution of particle diameters calculated from AFM images. **C)** *left:* Topography of a representative Ico8 particle imaged by conventional AFM; *middle and right*: images of Ico8 particles obtained by scanning PiFM at 1666 cm^-1^ **D)** IR spectrum of an individual Ico8 particle.

Further images and Infrared (IR) spectra for individual Ico8 particles were measured using IR-photo-induced force microscopy IR-PiFM (Fig. 3C,D). This technique records the vibrational spectrum of the material located in close proximity of an AFM tip operated in dynamic mode (26, 27). IR spectra recorded from individual particles showed characteristic amide-I and amide-II bands, confirming that the particles consist of proteins (Fig. 3D). Images obtained using the amide I band at 1666 cm^-1^ exhibited facets associated with the particles that are consistent with the formation of icosahedral cages. Interestingly, the particles appeared spherical under conventional AFM analysis, even when using a CNT tip.

### Cryo-electron microscopy of Ico8

Further structural information on the cages formed by Ico8, was obtained by cryo-EM single particle analysis. A total of 18,707 particles were excised then subjected to reference-free classification and averaging using the program RELION (28). 2D class-averaged cryo-EM images (Fig. S3) indicate Ico8 forms hollow cages that range in diameter from 20 to 30 nm, with an average diameter ~25 nm (Fig. 4A,B). Unexpectedly, two shells of density were observed in all the class averages, with most classes exhibiting short, inward-pointing features that appear to connect the inner and outer shells at multiple sites. The thickness and gap between the two shells appeared independent of the sizes of these cages (Fig S4). Interestingly, these inward-pointing features seem compatible with the size of the pentameric coiled coil domain (Fig. 4B).

**Figure 4.**
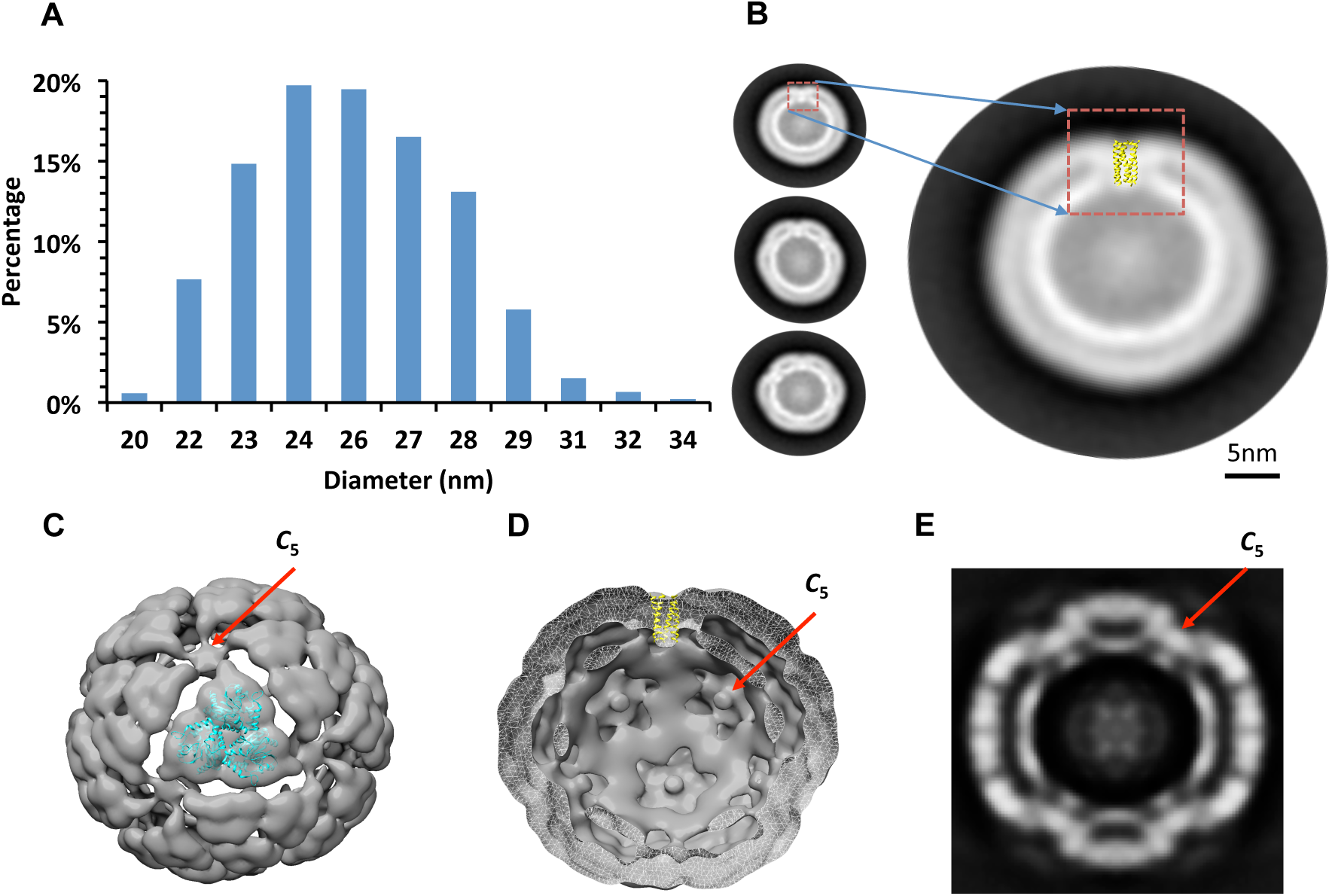
Cryo EM analysis of Ico8 cages. **A)** Size distribution of Ico8 cage measured from 2D class averages. **B)** Representative reference free 2D class averaged images of Ico8 cages illustrating the double-walled appearance of the cages and regularly spaced constrictions between the inner and outer walls that likely represent the coiled coil domain. The expanded view of one class averaged image shows coiled coil domain (yellow) superimposed on the density attributed the coiled coil. **C, D)** Cross-sections through the 3D electron density map of Ico8, with the crystal structures of TriEst **(C)** and the pentameric coiled coil **(D)** docked. **E)** Cross-section view of density map perpendicular to 2-fold axis. Red arrows indicate the 5-fold symmetry element specified by the coiled coil.

We then selected 10,363 particle projections, representing the average size of the protein cage, to calculate a low-resolution 3D cryo-EM reconstruction of Ico8 using the program cisTEM (29). Icosahedral symmetry was enforced during the refinement and the final reconstruction indicated a resolution of 12 Å. The symmetrically reconstructed electron density map clearly reveals the arrangement of esterase trimers that surround a region of inward-pointing density attributable to the 5-fold symmetric coiled coil domain (Figure 4C-E). This is consistent with inward-pointing density seen in the 2D-class averages. The orientation of the coiled coil, inward or outward, was not specified in the design and it is possible that there is a minor population of coiled coils that point outwards but are not resolved by cryo-EM. The inner shell of electron density is also clearly evident in the reconstruction.

It is evident that the electron density poorly accommodates the structure of TriEst, as the density is thinner and more spread out than expected (Fig. 4C). This is attributable to the flexible nature of the protein cage, which leads to smearing out of the averaged density. However, although of low resolution, the structure could be reproducibly reconstructed from the data. Furthermore, the icosahedral features of the cage were largely maintained when the symmetry used in the reconstruction was reduced to C_5_, providing confidence that the icosahedral geometry is correct.

### Ico8 encapsulates DNA

As noted above, we were surprised that the both 2D class averages and the 3D reconstruction identified an inner shell of strongly scattering electron density within the cages. Noting that nucleic acids scatter more strongly than proteins in cryo-EM images, and that thickness of the inner wall is similar to the diameter of a DNA double helix, we decided to investigate whether nucleic acids may be encapsulated within the cages; interestingly, this appears to be the case.

Protein cages were extracted with phenol:chloroform to remove the protein and the nucleic acids analyzed by electrophoresis, either before or after digestion with RNase A or DNase I. The recovered nucleic acids fragments were approximately 300 – 500 bp in size and could be digested by DNase I treatment (but not RNase A treatment) indicating that they comprise DNA (Fig S4). Quantification of the nucleic acid and protein content of the cages, determined by agarose electrophoresis and SDS-PAGE respectively, indicated that the nucleic acid component comprises ~30 % by weight of the purified Ico8 cages. We note that the cages are certainly large enough to accommodate this amount of DNA; indeed, viral capsids of similar diameters can encapsulate single-stranded genomes of 4 – 5 kbp (30). Therefore, we ascribe the inner “shell” of the cages to DNA that is non-specifically associated with the inner face of the protein cage.

### Stability of Ico8

Many natural protein cages, e.g. viral capsids, exhibit high stability towards unfolding. We therefore examined how assembly of the TriEst building block into an icosahedral cage affected its stability and catalytic activity. Circular dichroism (CD) was used to follow the thermally-induced unfolding of both Ico8 and TriEst by monitoring the decrease in ellipticity at 222 nm (Fig. 5A). Whereas TriEst irreversibly denatured between 70 and 80 °C, Ico8 remained folded at 98 °C (the highest temperature examined). The thermal stability of Ico8 was further investigated using differential scanning calorimetry (DSC), which permitted thermal unfolding of the protein to be studied at temperatures up to 120 °C at a pressure of 3 atm (Fig. S5). Again, TriEst underwent an irreversible exothermic transition between 70 °C and 80 °C, indicative of protein unfolding; however, Ico8 exhibited no thermal transitions up to 120 °C, implying that it remained folded. Consistent with this, no discernable change in morphology was apparent when the protein cages were imaged by EM after cooling (Fig. S6). The esterase building block retained full catalytic activity, determined with 2,4-dinitrophenol acetate as substrate, when assembled into Ico8 cages and, significantly, remained fully active after heating and cooling (Table S2). In contrast, as expected, TriEst was completely inactivated by heat treatment.

**Figure 5.**
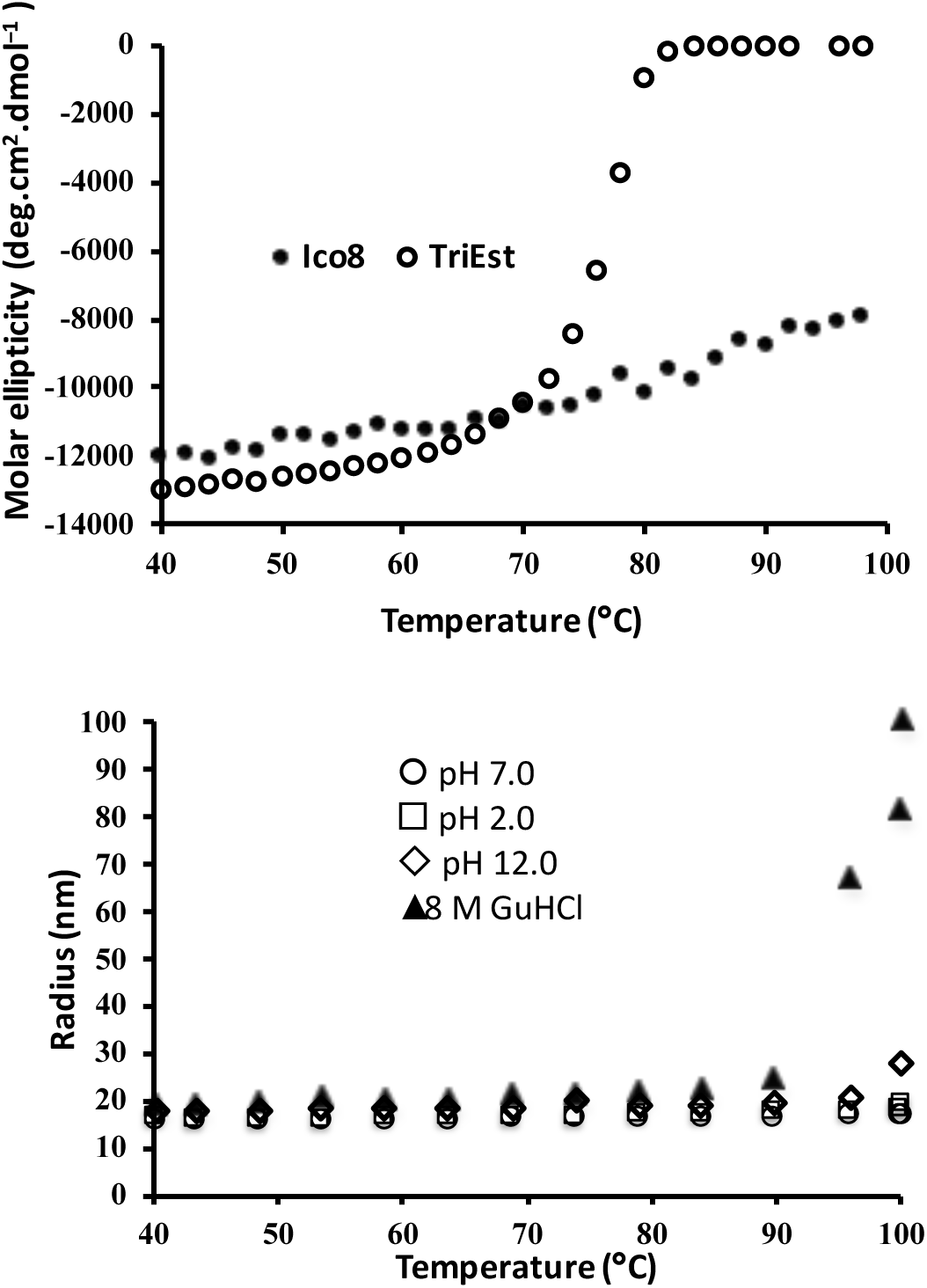
Thermal and chemical stability of Ico8 protein cage. **A)** The thermally-induced unfolding of TriEst and Ico8 monitored by changes in molar ellipticity at 222 nm. **B)** Thermally-induced unfolding of Ico8 monitored by light scattering as a function of temperature at both high and low pH and in 8 M GuHCl.

The cages formed by Ico8 also proved extremely stable towards chemical denaturation. We compared the stability of Ico8 and TriEst at pH 2.0 and pH 12.0. At low pH the TriEst building block was unfolded and/or aggregated. However, both CD spectroscopy and DLS measurements indicated that the cages formed by Ico8 remain intact. Similarly, Ico8 remained folded even in the presence of 8 M GuHCl, whereas TriEst was irreversibly unfolded by ~ 1.5 M GuHCl (Fig. S7). We extended these measurements by using temperature-dependent DLS to examine the thermal stability of Ico8 at pH 2.0, at pH 12.0, and in 8 M GuHCl (Fig. 5B). The number-averaged diameter distribution of particles recorded by DLS provides a sensitive measure of protein aggregation, which is indicative of partial unfolding. At pH 2.0, Ico8 showed no evidence for unfolding/aggregation at temperatures up to 98 °C. At pH 12.0, or in 8 M GuHCl, an increase in particle diameter, indicative of aggregation, was only evident when the protein samples were heated to temperatures above 90 °C. Again, no discernable change in morphology was apparent when the protein cages were imaged by negative stain EM after treatment with denaturants. The only exception was samples heated in 8 M GuHCl, for which some aggregation was evident by EM (Fig. S6). Interestingly, when the cages heated in GuHCl were re-imaged after storing at room temperature for 2 weeks, the aggregated material was no longer evident although the cages appeared more heterogeneous in size. (Fig S6).

## Discussion

Using a simple, symmetry-based approach, we demonstrate the assembly from “off-the-shelf” components of a nano-scale protein container with a size comparable to a viral capsid. Remarkably, the combination of a C_3_-symmetric building protein block and a C_5_-symmetric coiled coil, appears to be sufficient to assemble an icosahedral protein cage. As with our previous tetrahedral and octahedral cage designs (17, 19), it was necessary to optimize the linker length to achieve efficient assembly, but this required screening only a small number of constructs to identify a successful design. Importantly, the assembly process does not compromise the catalytic activity of the enzyme, which is actually higher for Ico8 than the esterase from which Ico8 is assembled (Table S2). Thus, our design approach appears to be both an efficient and general method to assembling proteins that requires minimal computational modeling to implement. The relative ease with which tetrahedral, octahedral and icosahedral protein cages are formed suggests that natural protein cages may perhaps have evolved through a similar route involving gene fusion of C_3_-symmetric proteins with C_3_-, C_4_- or C_5_-symmetric proteins. If the cages conferred an advantage on the organisms in which they arose, then further rounds of natural selection may have led to the development of more rigid cages with extensive protein interfaces.

The size distribution of the Ico8 cages is broader than that seen for the octahedral and tetrahedral cages we described previously (17, 19). This probably reflects the flexibility of the cages that arises from the flexible 8-Gly linker connecting the TriEst and coiled coil domains and the fact this icosahedral cage is much larger than the octahedral and tetrahedral cages. This flexibility is evident in the electron density of the reconstructed structure obtained by cryo-EM, in contrast, and perhaps unsurprisingly, better-defined cryo-EM reconstructions were obtained for the smaller tetrahedral and octahedral cages assembled with coiled coils. Further evidence for Ico8 forming 60-subunit icosahedral structures comes from individual EM and PiFM images where hexagonal features consistent with icosahedral geometry are also clearly evident.

An interesting feature of the cages formed by Ico8 is their extremely high thermal and chemical stability. Whereas TriEst is irreversibly unfolded at 75 °C, and in the presence of 1.5 M GuHCl, the Ico8 cages remain intact at 120 °C and in 8 M GuHCl. This places Ico8 among some of the most thermostable proteins so far described, either natural or designed. The enzymatic activity of the cages is not affected by exposure to these harsh conditions, which is clearly a valuable feature for many biotechnological applications. An icosahedral protein cage, thermostable to 90 °C, which was engineered by designing new subunit interfaces, was recently reported by Baker and co-workers (3). However, in this case the protein building block was obtained from a hyperthermophile and so was intrinsically thermostable. As shown in Fig. 5A, TriEst is not itself very thermostable, posing a question as to the origin of the remarkable stability exhibited by Ico8. This may partly derive from the stability of the coiled coil domain, which was previously shown to withstand thermal unfolding (25). Further, cooperative effects that arise from the increased number of inter-subunit interactions in the assembled cage, also may contribute to Ico8’s stability. Similar cooperative interactions between capsid proteins are suggested to the high stability of many viral capsids (31-33). The high rigidity of viral capsids has also been cited as a source of their stability (31-33); however, our results suggest that, contrary to prevailing assumptions, protein rigidity is not a prerequisite for assembling extremely stable protein cages.

An unanticipated feature of the cages formed by Ico8 is that they appear to encapsulate nucleic acids, most likely DNA. When visualized by cryo-EM the nucleic acid appears to form a separate shell with in the protein cage. This was unexpected because neither TriEst, nor the coiled coil binds nucleic acids, and nucleic acids did not bind to either of the previously constructed octahedral or tetrahedral cages. In this respect, nucleic acid encapsulation may be considered a truly emergent property of Ico8 because it is not a property of the protein components and was not intentionally designed into the cage. This observation intriguingly hints at a mechanism whereby viruses could have evolved, de novo, from cellular components through the adventitious encapsulation of nucleic acids.

## Materials and Methods

### Construction of genes encoding fusion proteins

Codon-optimized genes ligated into the expression vector pET28b were either commercially synthesized or derived from the other constructs using standard techniques. The sequences of the proteins are included in Table S1.

### Protein expression and purification

Expression constructs were transformed into *E. coli* BL21(DE3) cells and protein expression induced by IPTG using standard methods as described previously (17, 19). For initial characterization, proteins were purified by standard methods using an N-terminal 6-His tag to facilitate affinity chromatography on a Ni-NTA column as described previously (17, 19). For further characterization of Ico8, the protein was purified as follows: ~5 g of cells was resuspended on ice in 45 mL of 50 mM HEPES buffer, pH 7.5 containing 300 mM ammonium acetate, 50 mM imidazole, 1.5 M urea and 5% (v/v) glycerol; a protease inhibitor tablet (benzonase) and 50 mg of lysozyme were added and the suspension gently shaken for 20 min. The cells were lysed by sonication and the lysate clarified by centrifugation for 30 min at 40,000 *g*. The supernatant was passed through a 5 mL Ni-NTA column at 0.25 mL/min and the flow-through heated at 70 °C for 75 min. Precipitated proteins were removed by centrifugation at 40,000 *g* for 30 min and the supernatant dialyzed at 4 °C against 20 mM, pH 7.5, HEPES, 2 mM EDTA, 100 mM ammonium acetate for one week. Dialyzed samples were further purified by 2 rounds of SEC using a Superose-6 300/10 column equilibrated in the same buffer (flow rate 0.3 mL/min). Fractions containing Ico8 were pooled, concentrated by ultrafiltration and rechromatographed on the same column. Samples were then concentrated and stored at room temperature.

### Enzyme Activity

The esterase activity of Ico8 and Tri-Est were assayed using 2,4-dinitrophenol acetate as the substrate and measuring the increase in absorbance at 405 nm, as previously described (17).

### Nucleic acid analysis

Nucleic acids were purified from protein samples by phenol:chloroform extraction using standard methods. The presence of DNA, rather than RNA, was confirmed by digestion with either DNAase I (RNAase free) or RNAase, followed by gel electrophoresis. 500 ng of phenol:chloroform extracted nucleic acids were incubated with either 1 unit of DNAase I or RNAase A for 30 min at 25 °C in the manufacturers’ buffer. The amount of DNA was quantified by analysis of agarose gels and comparison to DNA standards using the program Image J.

### Electron Microscopy

Proteins (0.03 – 0.1 mg/mL) were fixed on Formvar/Carbon 400 Mesh, Cu grids using conventional procedures and staining with uranyl formate. Imaging was performed at room temperature with JEOL 1500 electron microscope equipped with tungsten filament, XR401 high sensitivity CMOS camera and operated at 90 keV. Samples for scanning transmission electron microscopy were prepared similarly. Images were acquired using a JEOL JEM-2100F transmission electron microscope (TEM) with a CEOS probe corrector. Microscopy was performed at 200 keV in scanning transmission microscopy (STEM) mode with the lens setting corrected by the Cs-corrector to produce a sub-Angstrom beam size. Both high-angle annular dark-field and bright-field images were acquired simultaneously. To better display the internal microstructures of the virus, fast Fourier transform image filtering was performed for bright field images.

### Dynamic Light Scattering

DLS measurements were made using a DynaPro Nanosizer ZS instrument. Measurements were made at 25 °C with the refractive index and absorption coefficient for the particles were set at 1.45 and 0.001 respectively. Diameter distributions by intensity were recorded. Measurements were made in triplicate with each measurement comprising an average of 15 scans. Temperature-dependent DLS was performed using the same instrument. 10 μl samples with protein concentration 0.2 mg/ml were heated at a rate of 5 °C/min from 40 to 100 °C. The refractive index and absorption coefficient for the particles were set at 1.45 and 0.001 respectively, and the diameter distribution by intensity was recorded.

### Circular Dichroism (CD)

CD measurements were performed using Jasco J815 CD spectrometer. Protein concentrations were between 0.5 – 0.2 mg/mL, (1 mm path length cuvette). Samples were heated from 40 °C to 98 °C at a rate of 1 °C /min.

### Differential Scanning Calorimetry (DSC)

DSC scans were performed using a TA instruments nanoDSC, with dialysis buffer was used as reference. ~ 0.6 mL of samples of protein, 0.3 - 0.4 mg/mL, were heated at 1 °C/min from 50 °C to 115 °C at 3 atm pressure and the changes in heat taken up by the samples recorded.

### Atomic force microscopy (AFM)

Samples for AFM analysis were first chemically cross-linking using the lysine-specific reagent bis(sulfosuccinimidyl)suberate (Thermo Fisher) as previously described (17) to achieve ~ 50% cross-linking as determined by SDS-PAGE. The protein solution (20 μL ~ 0.2 mg/mL) was spin-coated at 3500 rpm onto a freshly cleaved mica surface and further rinsed with 40 μL milli-Q water while spinning to remove salt from the surface. AFM imaging was carried out under AC mode using PicoPlus 5500 (Agilent Electronic Measurement, K-tek CNT tip) AFM system. A total of 20 images containing ~1100 cages were analyzed by the automated particle detection function in SPIP (version 6.2.6, Image Metrology, 0.8 nm z-range cutoff threshold). The diameter was calculated from the measured volume using the icosahedral geometric relationship: 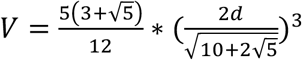, where V is the volume and d is the diagonal diameter.

The IR-PiFM measurements were performed using a Molecular Vista Inc, Vista-IR microscope. A QCL based light source illuminated the AFM probe (300 kHz NCHR-Au, NanoSensors AG) providing a spectral range from 770-1885 cm^-1^. The protein solution (20 uL, 0.2 mg/mL) was drop-cast onto freshly cleaved mica and air dried before imaging. The spectral acquisition time was 1.0 seconds with image acquisition time of 4.2 min at 256 × 256 pixels.

### Cryo-electron microscopy

Samples were concentrated to ~ 0.3 mg/ml and loaded onto glow-discharged QUANTIFOIL R2/2 200 mesh grids and flash-frozen using Vitrobot (FEI Mark IV). The samples were visualized at liquid nitrogen temperature on a Talos Arctica electron microscope (FEI) operating at 200 kV. Cryo-EM images were recorded at a nominal magnification of ×34014 using a K2 Summit direct electron detector (Gatan Inc.) in counted mode.

A total of 560 micrographs were taken using automation software SerialEM (34), Dose-fractionated image Images were subjected to motion correction using MotionCor2 (35), and binned by 2 resulting in a sampling of 2.94 Å per pixel for particle picking and processing. 18,707 particles were selected automatically, then subject to reference-free 2D classifications using RELION (28). Well-defined class average images were selected for further 3D reconstruction using cisTEM (29).

## Author contributions

A.S.C-D., J.C., D.B.N. and M.S. performed the experiments. A.S.C-D., J.C., D.B.N., M.S., M.M.B.H. and E.N.G.M. designed the experiments. A.S.C-D., J.C., D.B.N., M.S., M.M.B.H. and E.N.G.M. analyzed the data and wrote the manuscript.

## Acknowledgements

We thank Dr. Kai Chen, University of Michigan for help in acquiring STEM images. This work was supported in part by grants from the National Institutes of Health (GM 093088) and the Army Research Office (W911NF-11-1-0251) to E.N.G.M.

